# Unexpected contribution to the prevailing trend of positive results for 40 Hz light flicker

**DOI:** 10.1101/2023.10.27.564342

**Authors:** M. Carstensen, J. T. Pedersen, J. Carstensen

## Abstract

This comment addresses the potential influence of sample size on the conclusions drawn from a study investigating the beneficial effects of neuropathology in Alzheimer’s disease (AD). By examining the existing literature and employing statistical reasoning, we argue that an increase in sample size by factor of 2-4 would have led the authors to arrive at a different conclusion. Contrary to expectations, their findings unexpectedly contribute to the prevailing trend of positive results regarding the advantageous impacts of neuropathology in AD, with nine out of ten of their immunohistochemistry experiments showing a consistent ∼30 % reduction of amyloid plaque. We demonstrate that the quantity and quality of the data presented by Soula et al. 2023 do not support the paper’s conclusions on amyloid neuropathology. Based on a thorough statistical analysis of the available data, we therefore submit that given a larger sample size, the conclusion would have been positive towards possible improvements in neuropathology.

## Background

The stimulation of brain oscillations through non-invasive light treatment has garnered substantial attention in the study of neurological disorders, notably Alzheimer’s disease (AD). In the existing literature^2-6^, a wealth of evidence is provided, derived from both mouse and human studies, consistently supporting the positive effects of 40 Hz light flicker on neuropathology. Moreover, these findings extend across different modalities^7-11^, including but not limited to sound, magnetic stimulation, infrared stimulation, and electrical stimulation., illustrating a long record of positive results carried by the 40-Hz modulation frequency. Notably, within the same studies, multiple independent research teams have consistently demonstrated that applying 40-Hz (gamma) visual light stimulation induces gamma power in the brains of mice and humans. Recently, Soula and colleagues^**1**^ attempted to repeat the original finding by Iccarino et al.^2^ and concluded that there was no significant effect of 40-Hz light stimulation on the neuropathology of AD mouse models.

### Data re-analysis

Alzheimer’s pathologic change is mainly defined by amyloid through the 2018 NIA-AA Research Framework. Pathological studies using immunochemistry are typically performed on a wide range of Alzheimer’s mouse models to provide a surrogate measure for meaningful clinical changes, e.g., as done by Iccarino et al.^2^. In the current paper by Soula et al.^1^, a mix of APP/PS1 and 5XFAD mice has been used. These two mouse strains have been extensively characterised, and their temporal development of neuropathology is well understood.

No age-dependent increase in plaque density was observed for any of the mice (Extended Fig. 1, Soula et al.^1^). Age-dependent plaque increase has been reported in both APP/PS1 (dE9 Jax Stock No.004462)^12^ and 5xFAD (Tg6799 Jax Stock No. 034840) mice^13^. The APP/PS1 mice used in this study are expected to exhibit progressive increase in amyloid pathology from 6 to 12 month of age^12^, and the 5xFAD strain used is expected to exhibit plaque pathology already from the age of two month gradually exhibiting increase in plaque spreading and density until 16 months of age^13^. The expected pathological progression of both these mouse strains is well-reviewed in the AlzForum transgenic model database^14^. Soula et. al. reported a >1% increase in plaque area for the APP/PS1dE9 mice from 7 to 12 months of age (total area approximately 1.5 % at 12 months). Similarly, for the 5xFAD mice a 0.5 % increase in plaque area is observed from 4 to 7 months of age (total area 2 % at 7 months of age). This indicates a very small observed progress in pathology.

Soula et al.^1^ base their conclusions on a nonparametric two-tailed Wilcoxon’s rank-sum test of control versus treatment groups. Differences between male and female mice, as well as variation among cohorts with different strains and age, were not considered, even though including such variations can have a considerable impact on the statistical inference. Unbalanced design, as in Soula et al.^1^ with uneven numbers of male and female mice in control and treatment groups, can bias results. Additionally, the variation in the temporal development of neuropathology has not been taken into account for the acute experiment. The chronic experiment only used one cohort (5xFAD 7-month mice), and therefore, more general inferences about the potential treatment effect cannot be drawn.

Through a reanalysis of the data with a more appropriate statistical model, we found consistent responses across most mouse brain parameters (Table 1). Nine out of 10 analyses showed reduction of plaque area and amyloid beta in mice exposed to 40-Hz light treatment, ranging from -19% to -41% with an average of -31%. Only the plaque area in the hippocampus during the chronic experiment increased by 14%. However, Bradford concentrations for A*β*42 in V1 and for A*β*40 in both hippocampus and V1 were significantly reduced (*P*<0.05). Half of the brain parameters exhibited significant differences between male and female mice, whereas the treatment effect did not change with the sex (all *P*>0.05). The power for testing the treatment effect in the two experiments was generally low (3-83%) despite the considerable effect sizes. Consequently, the lack of significance^1^ is primarily due to insufficient sample sizes (n=42 and n=23 mice for acute and chronic experiment, respectively).

**Table 1:** Reanalysis and power analysis of the mouse brain parameters (PA = Plaque Area, Ab peptide levels) in cortex, hippocampus (HC) and V1 from Soula et al.^1^ . Estimated effect sizes from 40-Hz treatment and P-values for all fixed effects (t=treatment, s=sex, t×s=interaction term) are from mixed model (significant effects at 95% confidence level are highlighted in bold). The power for testing the treatment effect is based on the Soula et al. sampling design, whereas the power simulation describes the minimum number of mice and the relative increase in sampling effort to be able to identify a 30% treatment effect with 80% probability.

| Experiment | Reference | Parameter | Effect sizes | Test for fixed effects |  |  | Power test treatment | Power simulation |  |
| --- | --- | --- | --- | --- | --- | --- | --- | --- | --- |
|  |  |  |  | *P(t)* | *P(s)* | *P(txs)* |  | Min n | Factor |
| Acute | Fig. 1d | PA Cortex | -35% | 0.3591 | **0.0050** | 0.8352 | 18% | 136 | 3.2 |
|  | Fig. 1e | PA HC | -30% | 0.5924 | 0.2590 | 0.6665 | 3% | 344 | 8.0 |
|  | Fig. 1f | PA V1 | -32% | 0.3271 | **0.0004** | 0.7778 | 7% | 176 | 4.1 |
| Chronic | Fig. 2a | PA Cortex | -19% | 0.2005 | 0.5617 | 0.1582 | 23% | 40 | 1.7 |
|  | Fig. 2a | PA HC | 14% | 0.4314 | **0.0299** | 0.7881 | 10% | 52 | 2.3 |
|  | Fig. 2a | PA V1 | -29% | 0.1661 | 0.1856 | 0.9026 | 27% | 78 | 3.4 |
| | Fig. 2c | Aβ42 HC | -21% | 0.0532 | 0.5148 | 0.0958 | 53% | 24 | 1.0 |
| | Fig. 2c | Aβ42 V1 | -34% | **0.0180** | **<0.0001** | 0.8262 | 59% | 36 | 1.6 |
| | Fig. 2c | Aβ40 HC | -41% | **0.0455** | 0.3639 | 0.9812 | 54% | 80 | 3.5 |
| | Fig. 2c | Aβ40 V1 | -36% | **0.0067** | **0.0050** | 0.2056 | 83% | 30 | 1.3 |

Our power analysis for the 10 brain parameters suggested sample sizes ranging from 24 to 344 mice are sufficient for identifying an effect size of 30% with at least 80% probability (Table 1). This corresponds to a 2-4-fold increase of the existing sampling design for most parameters, except for plaque area in the hippocampus for the acute experiment. These results underline that the lack of significance most likely stems from insufficient sample sizes rather than the absence of an effect of 40 Hz light stimulation. We submit that the authors would have arrived at a different conclusion had the sample size been 2-4 times larger. Lack of statistical power is frequently the source of unsupported conclusions; a seminal meta-analysis of studies in neuroscience concluded that the median power of studies in the scientific literature is between 8 and 31 %^15^.

## METHODS SUMMARY

We reanalysed the data from the acute and chronic experiments (Fig. 1d-f and Fig. 2a,c in Soula et al.^1^) with a mixed model that included sex (s: male versus female) and treatment (t: 40-Hz treatment versus control) as well as their interaction (s×t) as fixed effects. Cohort (C) and the interaction between cohort and treatment (C×T) were included as random effects for the acute experiment, but not in the chronic experiment that used one cohort only. Random effects for higher order interactions with sex (C×S and C×S×T) were not included due to the low number of observations. All variables were square root transformed to homogenise variation. Effect sizes were calculated from marginal means of the treatment effect after applying a back transformation to the original scale.

Variances from the mixed model were used to simulate (1000 realisations) the acute experiments with increasing number of cohorts (n=4 mice per cohort) and the chronic experiment with increasing number of mice using an average effect size of 30%. The percentage of significant outcomes at 95% confidence level was calculated as the power of the test. Simulated sample sizes achieving 80% power were reported. The power of the performed experiments was also calculated using estimated effect sizes.

## Supporting information

Supplement 1 Data analysis script

## Supplementary material files

SAS script files used for this analysis

Power analysis for brain study.sas

Brain stimulation analysis.sas

## Notes

https://www.nature.com/articles/s41593-023-01270-2

https://www.nature.com/articles/s41593-023-01270-2#Sec23

https://www.nature.com/articles/s41593-023-01270-2#Sec22

### Competing Interest Statement

MC is a fulltime employee and shareholder in Optoceutics ApS. JTP is a shareholder and independent consultant to OptoCutics ApS. JC declares no competing interests

