## Supplement 1 Data analysis script for "Unexpected contribution to the prevailing trend of positive results for 40 Hz light flicker"

```
/* Reanalysis of Figure 1 and Figure 2 data from Soula et al in Nature
Neuroscience */
/* Reading in brain data from excel files provided by Marisol Soula */
PROC IMPORT OUT=Fig1d_control
            DATAFILE= "F:\Temp\Marcus Carstensen\data
download\Soula_SourceData_Fig1_index.xlsx"
             DBMS=XLSX REPLACE;
     RANGE="d$A1:C21";
     GETNAMES=YES;
     MIXED=NO;
RUN;
PROC IMPORT OUT=Fig1d_treat
            DATAFILE= "F:\Temp\Marcus Carstensen\data
download\Soula_SourceData_Fig1_index.xlsx"
             DBMS=XLSX REPLACE;
     RANGE="d$d1:F23";
     GETNAMES=YES;
     MIXED=NO;
RUN;
data Fig1d;
   format brainpart $12. treatment $10.;
   set Fig1d_control(rename=(control=Plaque_area) in=ctrl)
Fig1d_treat(rename=(_40_Hz=Plaque_area) in=treat);
   brainpart='Cortex';
   if ctrl then treatment='Control';
   if treat then treatment='40 Hz';
   logPlaque_area=log(Plaque_area);
   sqrtPlaque_area=sqrt(Plaque_area);
   rename cohort_ID=cohort;
run;
PROC IMPORT OUT=Fig1e_control
            DATAFILE= "F:\Temp\Marcus Carstensen\data
download\Soula_SourceData_Fig1_index.xlsx"
             DBMS=XLSX REPLACE;
     RANGE="e$A1:C21";
     GETNAMES=YES;
     MIXED=NO;
RUN;
PROC IMPORT OUT=Fig1e_treat
            DATAFILE= "F:\Temp\Marcus Carstensen\data
download\Soula_SourceData_Fig1_index.xlsx"
             DBMS=XLSX REPLACE;
     RANGE="e$d1:F23";
     GETNAMES=YES;
     MIXED=NO;
RUN;
data Fig1e;
   format brainpart $12. treatment $10.;
   set Fig1e_control(rename=(control=Plaque_area) in=ctrl)
Fig1e_treat(rename=(_40_Hz=Plaque_area) in=treat);
   brainpart='Hippocampus';
   if ctrl then treatment='Control';
   if treat then treatment='40 Hz';
   logPlaque_area=log(Plaque_area);
   sqrtPlaque_area=sqrt(Plaque_area);
run;
PROC IMPORT OUT=Fig1f_control
```

```
            DATAFILE= "F:\Temp\Marcus Carstensen\data
download\Soula_SourceData_Fig1_index.xlsx"
             DBMS=XLSX REPLACE;
     RANGE="f$A1:C21";
     GETNAMES=YES;
     MIXED=NO;
RUN;
PROC IMPORT OUT=Fig1f_treat
            DATAFILE= "F:\Temp\Marcus Carstensen\data
download\Soula_SourceData_Fig1_index.xlsx"
             DBMS=XLSX REPLACE;
     RANGE="f$d1:F23";
     GETNAMES=YES;
     MIXED=NO;
RUN;
data Fig1f;
   format brainpart $12. treatment $10.;
   set Fig1f_control(rename=(control=Plaque_area) in=ctrl)
Fig1f_treat(rename=(_40_Hz=Plaque_area) in=treat);
   brainpart='V1';
   if sex="F'" then sex="'F'";
   if ctrl then treatment='Control';
   if treat then treatment='40 Hz';
   logPlaque_area=log(Plaque_area);
   sqrtPlaque_area=sqrt(Plaque_area);
run;
/* Analysis as in paper - Wilcoxon non-parametric test */
proc npar1way wilcoxon data=Fig1d;  * Same p-value as in paper for 2-
sided Normal approximation;
   class treatment;
   var Plaque_area;
   exact wilcoxon;
run;
proc npar1way wilcoxon data=Fig1e;  * Same p-value as in paper;
   class treatment;
   var Plaque_area;
   exact wilcoxon;
run;
proc npar1way wilcoxon  data=Fig1f;  * Same p-value as in paper;
   class treatment;
   var Plaque_area;
   exact wilcoxon;
run;
/* More correct analysis using mixed effects model - Neocortex */
/*proc mixed data=Fig1d covtest; * Full model;
   class treatment sex cohort;
   model sqrtPlaque_area = treatment sex treatment*sex;
   random cohort cohort*treatment cohort*sex cohort*treatment*sex;
   lsmeans treatment sex treatment*sex;
run;*/
proc mixed data=Fig1d covtest; * Reduced model - removing non-significant
random factors caused by data restrictions;
   class sex treatment cohort;
   model sqrtPlaque_area = treatment sex sex*treatment /residual
outp=res_Fig1d;
   random cohort cohort*treatment;
   lsmeans treatment sex sex*treatment;
   ODS output LSmeans=LSM_Fig1d;
```

```
run;
/* More correct analysis using mixed effects model - Hippocampus */
/*proc mixed data=Fig1e covtest; * Full model;
   class treatment sex cohort;
   model sqrtPlaque_area = treatment sex treatment*sex;
   random cohort cohort*treatment cohort*sex cohort*treatment*sex;
   lsmeans treatment sex treatment*sex;
run;*/
proc mixed data=Fig1e covtest; * Reduced model - removing non-significant
random factors caused by data restrictions;
   class sex treatment cohort;
   model sqrtPlaque_area = treatment sex sex*treatment /residual
outp=res_Fig1e;
   random cohort cohort*treatment;
   lsmeans treatment sex sex*treatment;
   ODS output LSmeans=LSM_Fig1e;
run;
/* More correct analysis using mixed effects model - V1 */
/*proc mixed data=Fig1f covtest; * Full model;
   class treatment sex cohort;
   model sqrtPlaque_area = treatment sex treatment*sex;
   random cohort cohort*treatment cohort*sex cohort*treatment*sex;
   lsmeans treatment sex treatment*sex;
run;*/
proc mixed data=Fig1f covtest; * Reduced model - removing non-significant
random factors caused by data restrictions;
   class sex treatment cohort;
   model sqrtPlaque_area = treatment sex sex*treatment /residual
outp=res_Fig1f;
   random cohort cohort*treatment;
   lsmeans treatment sex sex*treatment;
   ODS output LSmeans=LSM_Fig1f;
run;
/* Analysis of Figure 2 */
/* Reading in brain data from excel files provided by Marisol */
PROC IMPORT OUT=Fig2a_cortexcontrol
            DATAFILE= "F:\Temp\Marcus Carstensen\data
download\41593_2023_1270_MOESM6_ESM Fig 2.xlsx"
            DBMS=XLSX REPLACE;
     RANGE="a$A1:B12";
     GETNAMES=YES;
     MIXED=NO;
RUN;
PROC IMPORT OUT=Fig2a_cortextreat
            DATAFILE= "F:\Temp\Marcus Carstensen\data
download\41593_2023_1270_MOESM6_ESM Fig 2.xlsx"
            DBMS=XLSX REPLACE;
     RANGE="a$C1:D13";
     GETNAMES=YES;
     MIXED=NO;
RUN;
data Fig2a_cortex;
   format brainpart $12. treatment $10.;
   set Fig2a_cortexcontrol(rename=(Cortex_Control=Plaque_area) in=ctrl)
Fig2a_cortextreat(rename=(Cortex_40_Hz=Plaque_area) in=treat);
   brainpart='Cortex';
   cohort=10;
   if ctrl then treatment='Control';
```

```
   if treat then treatment='40 Hz';
   logPlaque_area=log(Plaque_area);
   sqrtPlaque_area=sqrt(Plaque_area);
run;
PROC IMPORT OUT=Fig2a_hippocontrol
            DATAFILE= "F:\Temp\Marcus Carstensen\data
download\41593_2023_1270_MOESM6_ESM Fig 2.xlsx"
            DBMS=XLSX REPLACE;
     RANGE="a$F1:G12";
     GETNAMES=YES;
     MIXED=NO;
RUN;
PROC IMPORT OUT=Fig2a_hippotreat
            DATAFILE= "F:\Temp\Marcus Carstensen\data
download\41593_2023_1270_MOESM6_ESM Fig 2.xlsx"
            DBMS=XLSX REPLACE;
     RANGE="a$H1:I13";
     GETNAMES=YES;
     MIXED=NO;
RUN;
data Fig2a_hippocampus;
   format brainpart $12. treatment $10.;
   set Fig2a_hippocontrol(rename=(Hippo_Control=Plaque_area) in=ctrl)
Fig2a_hippotreat(rename=(Hippo_40_Hz=Plaque_area) in=treat);
   brainpart='Hippocampus';
   cohort=10;
   if ctrl then treatment='Control';
   if treat then treatment='40 Hz';
   logPlaque_area=log(Plaque_area);
   sqrtPlaque_area=sqrt(Plaque_area);
run;
PROC IMPORT OUT=Fig2a_V1control
            DATAFILE= "F:\Temp\Marcus Carstensen\data
download\41593_2023_1270_MOESM6_ESM Fig 2.xlsx"
            DBMS=XLSX REPLACE;
     RANGE="a$K1:L12";
     GETNAMES=YES;
     MIXED=NO;
RUN;
PROC IMPORT OUT=Fig2a_V1treat
            DATAFILE= "F:\Temp\Marcus Carstensen\data
download\41593_2023_1270_MOESM6_ESM Fig 2.xlsx"
            DBMS=XLSX REPLACE;
     RANGE="a$M1:N13";
     GETNAMES=YES;
     MIXED=NO;
RUN;
data Fig2a_V1;
   format brainpart $12. treatment $10.;
   set Fig2a_V1control(rename=(V1_Control=Plaque_area) in=ctrl)
Fig2a_V1treat(rename=(V1_40_Hz=Plaque_area) in=treat);
   brainpart='V1';
   cohort=10;
   if ctrl then treatment='Control';
   if treat then treatment='40 Hz';
   logPlaque_area=log(Plaque_area);
   sqrtPlaque_area=sqrt(Plaque_area);
run;
```

```
PROC IMPORT OUT=Fig2c_AB42Hippocontrol
           DATAFILE= "F:\Temp\Marcus Carstensen\data
download\41593_2023_1270_MOESM6_ESM Fig 2.xlsx"
            DBMS=XLSX REPLACE;
     RANGE="c$A2:B13";
     GETNAMES=YES;
     MIXED=NO;
RUN;
PROC IMPORT OUT=Fig2c_AB42Hippotreat
           DATAFILE= "F:\Temp\Marcus Carstensen\data
download\41593_2023_1270_MOESM6_ESM Fig 2.xlsx"
            DBMS=XLSX REPLACE;
     RANGE="c$D2:E14";
     GETNAMES=YES;
     MIXED=NO;
RUN;
data Fig2c_AB42Hippo;
   format brainpart $12. treatment $10.;
   set Fig2c_AB42Hippocontrol(rename=(Hippo_Control=Plaque_area) in=ctrl)
Fig2c_AB42Hippotreat(rename=(Hippo_40_Hz=Plaque_area) in=treat);
   brainpart='Hippocampus';
   cohort=10;
   if ctrl then treatment='Control';
   if treat then treatment='40 Hz';
   logPlaque_area=log(Plaque_area);
   sqrtPlaque_area=sqrt(Plaque_area);
run;
PROC IMPORT OUT=Fig2c_AB42V1control
           DATAFILE= "F:\Temp\Marcus Carstensen\data
download\41593_2023_1270_MOESM6_ESM Fig 2.xlsx"
            DBMS=XLSX REPLACE;
     RANGE="c$H2:I13";
     GETNAMES=YES;
     MIXED=NO;
RUN;
PROC IMPORT OUT=Fig2c_AB42V1treat
           DATAFILE= "F:\Temp\Marcus Carstensen\data
download\41593_2023_1270_MOESM6_ESM Fig 2.xlsx"
            DBMS=XLSX REPLACE;
     RANGE="c$K2:L14";
     GETNAMES=YES;
     MIXED=NO;
RUN;
data Fig2c_AB42V1;
   format brainpart $12. treatment $10.;
   set Fig2c_AB42V1control(rename=(V1_Control=Plaque_area) in=ctrl)
Fig2c_AB42V1treat(rename=(V1_40_Hz=Plaque_area) in=treat);
   brainpart='V1';
   cohort=10;
   if ctrl then treatment='Control';
   if treat then treatment='40 Hz';
   logPlaque_area=log(Plaque_area);
   sqrtPlaque_area=sqrt(Plaque_area);
run;
PROC IMPORT OUT=Fig2c_AB40Hippocontrol
           DATAFILE= "F:\Temp\Marcus Carstensen\data
download\41593_2023_1270_MOESM6_ESM Fig 2.xlsx"
            DBMS=XLSX REPLACE;
```

```
     RANGE="c$A21:B32";
     GETNAMES=YES;
     MIXED=NO;
RUN;
PROC IMPORT OUT=Fig2c_AB40Hippotreat
           DATAFILE= "F:\Temp\Marcus Carstensen\data
download\41593_2023_1270_MOESM6_ESM Fig 2.xlsx"
            DBMS=XLSX REPLACE;
     RANGE="c$D21:E33";
     GETNAMES=YES;
     MIXED=NO;
RUN;
data Fig2c_AB40Hippo;
   format brainpart $12. treatment $10.;
   set Fig2c_AB40Hippocontrol(rename=(Hippo_Control=Plaque_area) in=ctrl)
Fig2c_AB40Hippotreat(rename=(Hippo_40_Hz=Plaque_area) in=treat);
   brainpart='Hippocampus';
   cohort=10;
   if ctrl then treatment='Control';
   if treat then treatment='40 Hz';
   logPlaque_area=log(Plaque_area);
   sqrtPlaque_area=sqrt(Plaque_area);
run;
PROC IMPORT OUT=Fig2c_AB40V1control
           DATAFILE= "F:\Temp\Marcus Carstensen\data
download\41593_2023_1270_MOESM6_ESM Fig 2.xlsx"
            DBMS=XLSX REPLACE;
     RANGE="c$H21:I32";
     GETNAMES=YES;
     MIXED=NO;
RUN;
PROC IMPORT OUT=Fig2c_AB40V1treat
           DATAFILE= "F:\Temp\Marcus Carstensen\data
download\41593_2023_1270_MOESM6_ESM Fig 2.xlsx"
            DBMS=XLSX REPLACE;
     RANGE="c$K21:L33";
     GETNAMES=YES;
     MIXED=NO;
RUN;
data Fig2c_AB40V1;
   format brainpart $12. treatment $10.;
   set Fig2c_AB40V1control(rename=(V1_Control=Plaque_area) in=ctrl)
Fig2c_AB40V1treat(rename=(V1_40_Hz=Plaque_area) in=treat);
   brainpart='V1';
   cohort=10;
   if ctrl then treatment='Control';
   if treat then treatment='40 Hz';
   logPlaque_area=log(Plaque_area);
   sqrtPlaque_area=sqrt(Plaque_area);
run;
/* Analysis of Fig 2a */
proc npar1way wilcoxon  data=Fig2a_cortex;  * Same p-value as in paper;
   class treatment;
   var Plaque_area;
   exact wilcoxon;
run;
proc npar1way wilcoxon  data=Fig2a_hippocampus;  * Same p-value as in
paper;
```

```
   class treatment;
   var Plaque_area;
   exact wilcoxon;
run;
proc npar1way wilcoxon  data=Fig2a_V1;  * Same p-value as in paper;
   class treatment;
   var Plaque_area;
   exact wilcoxon;
run;
proc mixed data=Fig2a_cortex covtest;
   class sex treatment;
   model sqrtPlaque_area = treatment sex sex*treatment /residual
outp=res_Fig2a_cortex;
   lsmeans treatment sex sex*treatment;
   ODS output LSmeans=LSM_Fig2a_cortex;
run;
proc mixed data=Fig2a_hippocampus covtest;
   class sex treatment;
   model sqrtPlaque_area = treatment sex sex*treatment /residual
outp=res_Fig2a_hippocampus;
   lsmeans treatment sex sex*treatment;
   ODS output LSmeans=LSM_Fig2a_hippocampus;
run;
proc mixed data=Fig2a_V1 covtest;
   class sex treatment;
   model sqrtPlaque_area = treatment sex sex*treatment /residual
outp=res_Fig2a_V1;
   lsmeans treatment sex sex*treatment;
   ODS output LSmeans=LSM_Fig2a_V1;
run;
/* Analysis of Fig 2c */
proc npar1way wilcoxon  data=Fig2c_AB42Hippo;  * Not exactly same p-value
as in paper;
   class treatment;
   var Plaque_area;
   exact wilcoxon;
run;
proc npar1way wilcoxon  data=Fig2c_AB42V1;  * Not exactly same p-value as
in paper;
   class treatment;
   var Plaque_area;
   exact wilcoxon;
run;
proc npar1way wilcoxon  data=Fig2c_AB40Hippo;  * Not exactly same p-value
as in paper;
   class treatment;
   var Plaque_area;
   exact wilcoxon;
run;
proc npar1way wilcoxon  data=Fig2c_AB40V1;  * Not exactly same p-value as
in paper;
   class treatment;
   var Plaque_area;
   exact wilcoxon;
run;
/* Fig. 2c - mixed model */
proc mixed data=Fig2c_AB42Hippo covtest;
   class sex treatment;
```

```
   model sqrtPlaque_area = treatment sex sex*treatment /residual
outp=res_Fig2c_AB42Hippo;
   lsmeans treatment sex sex*treatment;
   ODS output LSmeans=LSM_Fig2c_AB42Hippo;
run;
proc mixed data=Fig2c_AB42V1 covtest;
   class sex treatment;
   model sqrtPlaque_area = treatment sex sex*treatment /residual
outp=res_Fig2c_AB42V1;
   lsmeans treatment sex sex*treatment;
   ODS output LSmeans=LSM_Fig2c_AB42V1;
run;
proc mixed data=Fig2c_AB40Hippo covtest;
   class sex treatment;
   model sqrtPlaque_area = treatment sex sex*treatment /residual
outp=res_Fig2c_AB40Hippo;
   lsmeans treatment sex sex*treatment;
   ODS output LSmeans=LSM_Fig2c_AB40Hippo;
run;
proc mixed data=Fig2c_AB40V1 covtest;
   class sex treatment;
   model sqrtPlaque_area = treatment sex sex*treatment /residual
outp=res_Fig2c_AB40V1;
   lsmeans treatment sex sex*treatment;
   ODS output LSmeans=LSM_Fig2c_AB40V1;
run;
/* Extended Fig. 2 analysis */
/* Fig. 2a */
proc npar1way wilcoxon  data=Fig2c_AB40Hippo;  * Not exactly same p-value
as in paper;
   class treatment;
   var Plaque_area;
   exact wilcoxon;
   where sex="'M'";
run;
proc npar1way wilcoxon  data=Fig2c_AB40Hippo;  * Not exactly same p-value
as in paper;
   class treatment;
   var Plaque_area;
   exact wilcoxon;
   where sex="'F'";
run;
/* Fig. 2b */
proc npar1way wilcoxon  data=Fig2c_AB40V1;  * Same p-value as in paper;
   class treatment;
   var Plaque_area;
   exact wilcoxon;
   where sex="'M'";
run;
proc npar1way wilcoxon  data=Fig2c_AB40V1;  * Same p-value as in paper;
   class treatment;
   var Plaque_area;
   exact wilcoxon;
   where sex="'F'";
run;
/* Fig. 2c */
proc npar1way wilcoxon  data=Fig2c_AB42Hippo;  * Not exactly same p-value
as in paper;
```

```
   class treatment;
   var Plaque_area;
   exact wilcoxon;
   where sex="'M'";
run;
proc npar1way wilcoxon  data=Fig2c_AB42Hippo;  * Not exactly same p-value
as in paper;
   class treatment;
   var Plaque_area;
   exact wilcoxon;
   where sex="'F'";
run;
/* Fig. 2d */
proc npar1way wilcoxon  data=Fig2c_AB42V1;  * Same p-value as in paper;
   class treatment;
   var Plaque_area;
   exact wilcoxon;
   where sex="'M'";
run;
proc npar1way wilcoxon  data=Fig2c_AB42V1;  * Same p-value as in paper;
   class treatment;
   var Plaque_area;
   exact wilcoxon;
   where sex="'F'";
run;


/*****************************************************************/
/* Combining LSmeans                                             */
/*****************************************************************/
data LSM_figs_all;
   format Figlabel $20.;
   set LSM_fig1d(in=fig1d) LSM_fig1e(in=fig1e) LSM_fig1f(in=fig1f)
LSM_fig1h(in=fig1h)
       LSM_fig2a_cortex(in=fig2a_cortex)
LSM_fig2a_hippocampus(in=fig2a_hippo) LSM_fig2a_V1(in=fig2a_V1)
LSM_fig2b_pcc(in=fig2b_pcc)
       LSM_fig2c_ab42hippo(in=fig2c_ab42hippo)
LSM_fig2c_ab42V1(in=fig2c_ab42V1) LSM_fig2c_ab40hippo(in=fig2c_ab40hippo)
LSM_fig2c_ab40V1(in=fig2c_ab40V1);
   if fig1d then Figlabel='Fig. 1d Cortex';
   if fig1e then Figlabel='Fig. 1e Hippocampus';
   if fig1f then Figlabel='Fig. 1f V1';
   if fig2a_cortex then Figlabel='Fig. 2a Cortex';
   if fig2a_hippo then Figlabel='Fig. 2a Hippocampus';
   if fig2a_V1 then Figlabel='Fig. 2a V1';
   if fig2c_ab42hippo then Figlabel='Fig. 2c AB42 Hippocampus';
   if fig2c_ab42V1 then Figlabel='Fig. 2c AB42 V1';
   if fig2c_ab40hippo then Figlabel='Fig. 2c AB40 Hippocampus';
   if fig2c_ab40V1 then Figlabel='Fig. 2c AB40 V1';
   median=estimate*estimate;
   upper=(estimate+stderr*tinv(.975,DF))*(estimate+stderr*tinv(.975,DF));
   lower=(estimate-stderr*tinv(.975,DF))*(estimate-stderr*tinv(.975,DF));
   if fig1h or fig2b_pcc then do;
      median=estimate; upper=estimate+stderr*tinv(.975,DF);
lower=estimate-stderr*tinv(.975,DF); end;
   plus=upper-median;
   minus=median-lower;
```

```
run;
proc sort data=LSM_figs_all; by sex treatment; run;
proc export data=LSM_figs_all outfile='F:\Temp\Marcus
Carstensen\SASoutput\Micestudy figs2.xlsx' DBMS=XLSX replace;
     SHEET="LSM_figs_all";
run;
```
